# Tumor Evolution Decoder (TED): Unveiling Tumor Evolution Based on Mutation Profiles of Subclones or Single Cells

**DOI:** 10.1101/633610

**Authors:** Yitan Zhu, Subhajit Sengupta, Lin Wei, Shengjie Yang, Yuan Ji

## Abstract

Cancer cells constantly evolve accumulating somatic mutations. To describe the tumor evolution process, we develop the Tumor Evolution Decoder (TED), a novel algorithm for constructing phylogenetic tree based on somatic mutation profiles of tumor subclones or single cells. TED takes a unique strategy that reduces the total number of duplicated mutations and dropout mutations in the tumor evolution process, which has not been explored by previous phylogenetic tree methods. TED allows multiple types of somatic mutations as input, such as point mutations, copy number alterations, gene fusion, and their combinations. Theoretical properties of TED are derived while its numerical performance is examined using simulated data. We applied TED to analyze single-cell sequencing data from an essential thrombocythemia tumor and a clear cell renal cell carcinoma, to investigate the ancestral relationships between tumor cells, and found genes related to disease initialization and development mutated in the early steps of evolution. We also applied TED to the subclones of a breast invasive carcinoma and provided important insights on the evolution and metastasis of the tumor.

## Introduction

All cancers go through evolution in which cells acquire mutations promoting tumor growth and malignancy^1-3^. Because somatic mutations do not necessarily occur in all the cells, tumors are heterogeneous containing cells of distinct genotypes. Most existing studies investigate tumor heterogeneity of single cells^4-7^ or subpopulations of cells called subclones^8-10^. While genetic profiling of single cells is the most precise means to assess intra-tumor heterogeneity, it still faces challenges such as amplification bias and relatively low genome coverage^11-12^. Also, it is currently difficult and costly to sequence a large number of cells and analyze the massive data. In contrast, DNA sequencing of a bulk sample remains the main approach to investigate intra-tumor diversity^11^. Bulk-sample DNA-sequencing data can be used to statistically infer the heterogeneous genomes of subclones, each of which consists of a group of cells sharing the same mutation profile. Various methods have been developed to perform such statistical analysis for identifying genotypes of subclones and their cellular proportions^13-16^.

Once the genotypes of single cells or subclones are known, either directly measured or statistically inferred, a key task is to uncover the evolutionary process of the distinct tumor genomes. Understanding tumor evolution helps elucidate how tumors arise, identify driver mutations, and investigate drug resistance mechanism^11^. Several existing tools have been developed to construct phylogenetic trees, such as MrBayes^17^, PAUP^18^, and PHYLIP^19^, some of which implement multiple methods. Most existing phylogeny methods can be grouped into two categories, *phenetic* methods and *cladistic* methods. Phenetic methods use distance measures to evaluate the differences between genomes and build a phylogenetic tree based on the distance matrix, such as UPGMA (unweighted paired group method with arithmetic mean)^20-21^, neighbor joining method^22-23^, minimum evolution method^24^, and Fitch-Margoliash method^25^. Cladistic methods are character-based methods and usually assume the genomes descend from a common ancestor and thus are closely related. Typical cladistic methods apply the idea of maximum parsimony that explains the observed data using a minimum number of evolution changes^26^, infer the phylogenetic tree through maximum likelihood estimation^27-28^, or take Bayesian approaches that assume a prior distribution of possible phylogenetic trees^17,28^.

We introduce the Tumor Evolution Decoder (TED), a computational approach specifically designed to reconstruct the evolution process of tumor based on the somatic mutation profiles of subclones or single cells. TED takes a novel and unique approach to reveal the ancestor-descendant relationship between tumor cells or subclones, which is different from the aforementioned methods. The following are several novel features of TED methodology.

1. TED constructs a phylogenetic tree of tumor genomes, each of which is represented by a mutation profile that can include multiple types of mutations, such as Single Nucleotide Variant (SNV), Copy Number Alteration (CNA), gene fusion, and others.
2. TED can infer unobserved genomes that are not included in the observed genomes (i.e. input data) but exist in the tumorgenesis evolution process.
3. TED is theoretically proven to construct a correct phylogenetic tree when there is no genotype calling error in data and there is no duplicated or dropout mutation in the evolution process to be recovered.

In addition, TED does not depend on the distance between genomes, which may lead to a wrong phylogeny if two less similar genomes actually descended from a more recent common ancestor. Furthermore, TED does not attempt to minimize the number of mutations in the evolution process, which enables TED to analyze samples possessing a large number of somatic mutations.

The remainder of the paper is organized as follows. The Methods Section introduces the properties and assumptions of tumor evolution process, develops the TED methodology that includes a phylogenetic tree construction algorithm and an edge pruning algorithm, and provides the theoretical foundation for TED. The Simulation Examples Section examines the performance of TED based on simulation datasets. The Results Section applies TED on single-cell sequencing datasets from an essential thrombocythemia tumor and a clear cell renal cell carcinoma, and one subclone dataset derived from a breast invasive carcinoma. The Conclusion and Discussion section summarizes the major contributions and findings and discusses potential future developments. We have prepared an open-source R package implementing the TED algorithm, which is accessible publicly at http://compgenome.org/ted/.

## Methods

Tumor somatic mutations refer to the changes in a tumor genome when compared to the corresponding normal genome. For simplicity, we assume the normal genome is the germline genome. Tables 1 and 2 illustrate how TED encodes SNV mutations and CNA mutations as mutation events, respectively. Multiple different situations of tumor genotypes and normal genotypes are illustrated. Some events, such as the mutation from homozygous wild type to homozygous variant (e.g. AA to BB in Table 1, where A and B are the wild type and variant, respectively), need to be encoded by more than one event. For the simplicity, we consider only SNV and CNA in this paper. But the proposed TED algorithm can be used for other mutation types as long as the mutation can be clearly defined by one or a set of mutation events. Notice that reverse mutation events, such as *CNA:* 3 → 2 (copy number changes from 3 to 2) and *CNA:* 2 → 3 (copy number changes from 2 to 3) at the same locus, are taken as different mutation events.

**Table 1.** Illustration of encoding a SNV mutation at a locus in both alleles.

| Normal Genotype \ Tumor Genotype | AA | AB |
| --- | --- | --- |
| AB | *SNV*: $0 \rightarrow 1$ | None |
| BB | *SNV*: $0 \rightarrow 1$, *SNV*: $1 \rightarrow 2$ | *SNV*: $1 \rightarrow 2$ |

A and B indicate the wild type and variant at the locus, respectively. AA, AB, and BB indicate homozygous wild type, heterozygous genotype, and homozygous variant, respectively.

**Table 2.** Illustration of encoding a CNA mutation at a locus.

| Normal Genotype \ Tumor Genotype | 1 copy | 2 copies | 3 copies |
| --- | --- | --- | --- |
| 1 copy | None | *CNA*: $2 \rightarrow 1$ | *CNA*: $3 \rightarrow 2$, *CNA*: $2 \rightarrow 1$ |
| 2 copies | CNA: 1 → 2 | None | CNA: 3 → 2 |
| 3 copies | CNA: 1 → 2, CNA: 2 → 3 | CNA: 2 → 3 | None |

### Assumptions and Properties of Tumor Evolution Process

We use *T* to denote the phylogenetic tree of a tumor evolution process (e.g. Fig. 1). Let *G*_1_, …, *G*_*N*_ denote the mutation profiles of *N* tumor genomes in *T*, where each 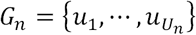 is a set of *U*_*n*_ mutation events each of which is denoted by *u*. For the simplicity, we call *G*_*n*_ a tumor genome and *G*_0_ is the normal genome, which is at the root of the phylogenetic tree and does not have any mutation event. By definition, *G*_0_ is an empty set denoted by *Φ*. Assume that no two tumor genomes are the same, i.e. *G*_*i*_ ≠ *G*_*j*_, ∀*i, j* ∈ {1, …, *N*} and *i* ≠ *j*. We call a branch starting from *G*_0_ (including all its descendants) a *lineage*. In Fig. 1 the phylogenetic tree is composed of two lineages. One starts with *G*_0_ → *G*_1_ and the other starts with *G*_0_ → *G*_2_. Within a lineage that includes *G*_*n*_, we use *A*(*G*_*n*_) and *D*(*G*_*n*_) to denote the set of genomes that are *ancestors* and *descendants* of *G*_*n*_, respectively. We use *E*_*n*_ to denote the *edge* that points to genome *G*_*n*_, which consists of the mutation events that are in *G*_*n*_ but not in its parent that is the direct ancestor of *G*_*n*_ and denoted by *P*(*G*_*n*_). Apparently, *P*(*G*_*n*_) ∈ *A*(*G*_*n*_). Mathematically, *E*_*n*_ = *G*_*n*_ − *P*(*G*_*n*_), including all the additional mutations that *P*(*G*_*n*_) needs to evolve to *G*_*n*_. The size of *E*_*n*_, i.e. |*E*_*n*_|, is called the length of edge *E*_*n*_. *Back mutations* (also called dropout mutations) of the same evolution step is denoted by *B*_*n*_. *B*_*n*_ = *P*(*G*_*n*_) − *G*_*n*_ is the set of mutation events in *P*(*G*_*n*_) but not kept in *G*_*n*_. Also, for any given edge *E*_*n*_, we can use 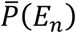 and 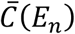 to denote the parent genome and the child genome on the edge, respectively. Apparently, 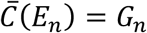 and 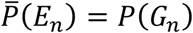.

**Figure 1.**
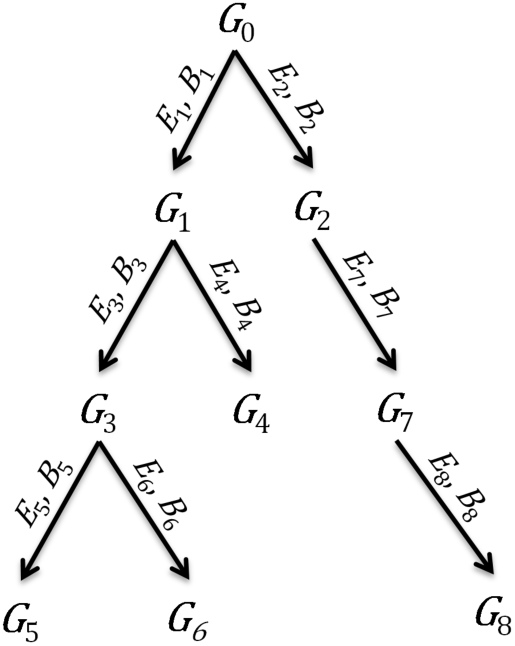
Illustration of a tumor evolution phylogenetic tree.

We make the following two assumptions about the tumor evolution process.

**Assumption 1:** No mutation event occurs twice during the evolution of a cancer case.

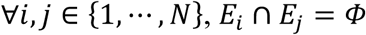

**Assumption 2:** No mutation event is ever lost, i.e. there is no back/dropout mutation.

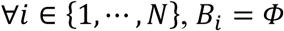

These two assumptions have been widely accepted in cancer evolution^11^ and constantly applied in the literature^1,11,29-30^, because both duplicated mutations and back mutations are rare events, requiring the same or reverse mutations to occur on the same locus of the genome twice during the evolutionary process^11,30^. Based on these two assumptions, we derive the following five properties of the phylogenetic tree and the genomes therein.

**Properties of a Phylogenetic Tree** that follows Assumptions 1 and 2

(1) ∀*i, j* ∈ {0,1, …, *N*}, *G*_*j*_ ∈ *A*(*G*_*i*_) ⟺ *G*_*j*_ ⊂ *G*_*i*_.

(2) ∀*i, j* ∈ {0,1, …, *N*}, *G*_*j*_ ∈ *D*(*G*_*i*_) ⟺ *G*_*j*_ ⊃ *G*_*i*_

(3) ∀*i, j, n* ∈ {0,1, …, *N*}, if *P*(*G*_*i*_) = *P*(*G*_*j*_) = *G*_*n*_, *G*_*i*_′ ∈ (*D*(*G*_*i*_) ∪ *G*_*i*_) and *G*_*j*_′ ∈ (*D*(*G*_*j*_) ∪ *G*_*j*_), then *G*_*i*_′ ⋂ *G*_*j*_′ = *G*_*n*_.

(4) ∀*i, j* ∈ {1, …, *N*}, if *G*_*i*_ ⊈ *G*_*j*_ and *G*_*i*_ ⊉ *G*_*j*_, then *G*_*i*_⋂*G*_*j*_ ∈ *A*(*G*_*i*_), *G*_*i*_⋂*G*_*j*_ ∈ *A*(*G*_*j*_), and *G*_*i*_⋂*G*_*j*_ is the closest to *G*_*i*_ and *G*_*j*_ and also the largest among all their common ancestors.

(5) ∀*i, j* ∈ {0,1, …, *N*}, *G*_*i*_⋂*G*_*j*_ ∈ {*G*_0_, *G*_1_, *G*_2_, … *G*_*N*_}

Properties 1-3 are quite obvious. We provide the proofs of Properties 4 and 5 in Supplementary Information Sections 1 and 2, respectively. Property 5 requires the intersection of any two genomes to be a genome in *T*, i.e. *G*_0_, *G*_1_, *G*_2_, … *G*_*N*_ are **closed under intersection**. We also call such a phylogenetic tree *T* closed under intersection. Property 4 and Property 5 together indicate the intersection of any two genomes in *T* must also appear in *T* as their closest and largest common ancestor.

If there is no error in genotype calling, we consider the data *noise-free*. However, real data always contain noise. A phylogenetic tree constructed based on real data can not always strictly satisfy the aforementioned assumptions and properties. Therefore, we define errors that measure the degree of violation. Specifically, *Type I error* counts the number of duplicated mutation events in all the edges *E*_1_, …, *E*_*N*_,

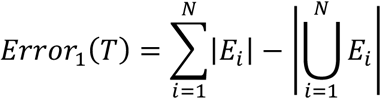

where |·| indicates the cardinality of a set. *Type II error* counts the total number of dropout mutations

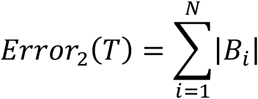

The summation of Type I and II errors gives the total error of constructing a phylogenetic tree,

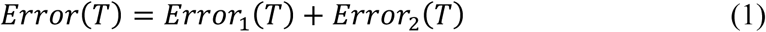

To measure how well an estimated phylogenetic tree (denoted by 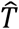) describes an evolution process (denoted by *T*), we introduce the concept of consistency as the following.

#### Definition 1 Consistency between Two Phylogenetic Trees

A phylogenetic tree 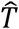 is consistent with another phylogenetic tree *T*, if (1) all genomes in 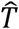 also appear in *T*, and (2) any two genomes in 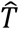 must correctly keep their relationship in *T*, which means if they have or have not an ancestor-descendant relationship in *T*, the same relationship also holds in 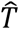.

Based on the definition, if 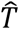 is consistent with *T*, there is no confliction between them in terms of topology. For a general case that may involve topology confliction, we can define a metric to measure the consistency level between two phylogenetic trees

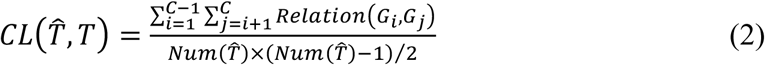

where *Num*(·) is a function counting the number of genomes in 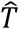 (including *G*_0_), *C* is the number of common genomes in 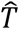 and *T* (including *G*_0_), and *Relation*(·,·) is an indicator function evaluating whether the relationship of two common genomes *G*_*i*_ and *G*_*j*_ changed between 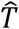 and *T*, given by

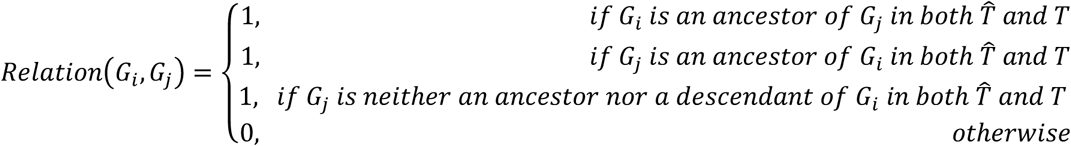

*CL*(·,·) basically counts the proportion of genome pairs with the same ancestor-descendant relationship in both 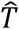 and *T*. Its value is between 0 and 1 and is 1 if and only if 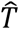 was consistent with *T*.

The error of an estimated phylogenetic tree is related to its level of consistency with the underlying evolution process to be estimated, as shown by the following theorem.

#### Theorem 1

Assume *T* is a phylogenetic tree satisfying Assumptions 1 and 2. Let *G*_0_, *G*_1_, …, *G*_*N*_ be the genomes in *T*. Let 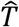 be another phylogenetic tree including *G*_0_ and a subset of {*G*_1_, …, *G*_*N*_}. Then, if 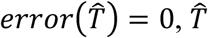 must be consistent with *T* and 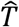 must be closed under intersection.

The proof of Theorem 1 is provided in Supplementary Information Section 3. Obviously, if 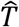 is consistent with *T*, 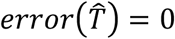 because *T* has 0 error. Therefore 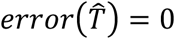 is the sufficient and necessary condition for 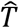 to be consistent with *T*. This naturally suggests that we should reduce the error of a phylogenetic tree when constructing it, and TED is such an algorithm.

### Phylogenetic Tree Construction Algorithm

Recall that the input data of TED are a group of tumor genomes, each consisting of a set of somatic mutation events. To start, TED takes two tumor genomes (e.g. *G*_*i*_ and *G*_*j*_ in Fig. 2) and generates an initial phylogenetic tree for the two tumor genomes and the normal genome, which can take one of the four forms in Fig. 2. Fig. 2d introduces a common parent of *G*_*i*_ and *G*_*j*_, i.e. *G*_*i*_ ∩ *G*_*j*_ ≠ *Φ*, which may be among the observed genomes or an unobserved genome involved in the tumor evolution process but not captured by the input data. Among the four possible initial trees, TED selects the tree with the minimum error. Actually, for two tumor genomes, one of the four possible initial trees must have 0 error (see the proof of Theorem 2).

**Figure 2.**
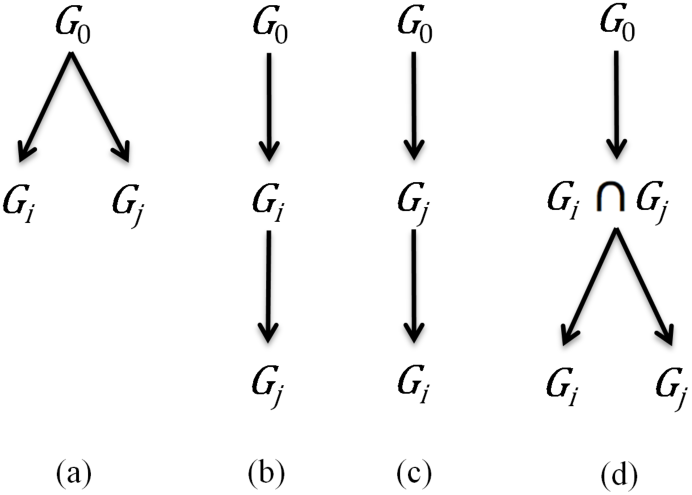
Four different schemes for generating an initial tree with *G*_*i*_ and *G*_*j*_

After constructing an initial tree, TED then proceeds by adding the other tumor genomes to the tree one by one. At each time, when growing the tree by including one more tumor genome, the algorithm considers all potential candidate tumor genomes, which are the observed genomes not included in the tree yet, and searches among all possible schemes for adding a candidate genome to the existing tree, to identify the tree with the minimum error. When adding a candidate genome (denoted by *G*_*m*_) to an existing tree, there are three possible types of schemes as shown in Fig. 3b-d. The first type of schemes adds *G*_*m*_ as a leaf child node to an existing node in the tree (Fig. 3b). The second type of schemes adds *G*_*m*_ as an intermediate node on an edge in the tree between a pair of parent and child nodes (Fig. 3c). In the third type of schemes (Fig. 3d), a latent intermediate node is added on an existing edge between a pair of parent and child nodes already in the tree and *G*_*m*_ is added as a leaf child node of the intermediate node. In general, this latent intermediate node is the intersection of *G*_*m*_ and the child node on the original edge (e.g. *G*_3_ in Fig. 3d). It may be observed in the input data or unobserved and not included in the input data, yet according to Property 4 it is a common ancestor of *G*_*m*_ and *G*_3_. For each type of schemes, all potential positions where the candidate genome can be added are considered for identifying the tree with the minimum error.

**Figure 3.**
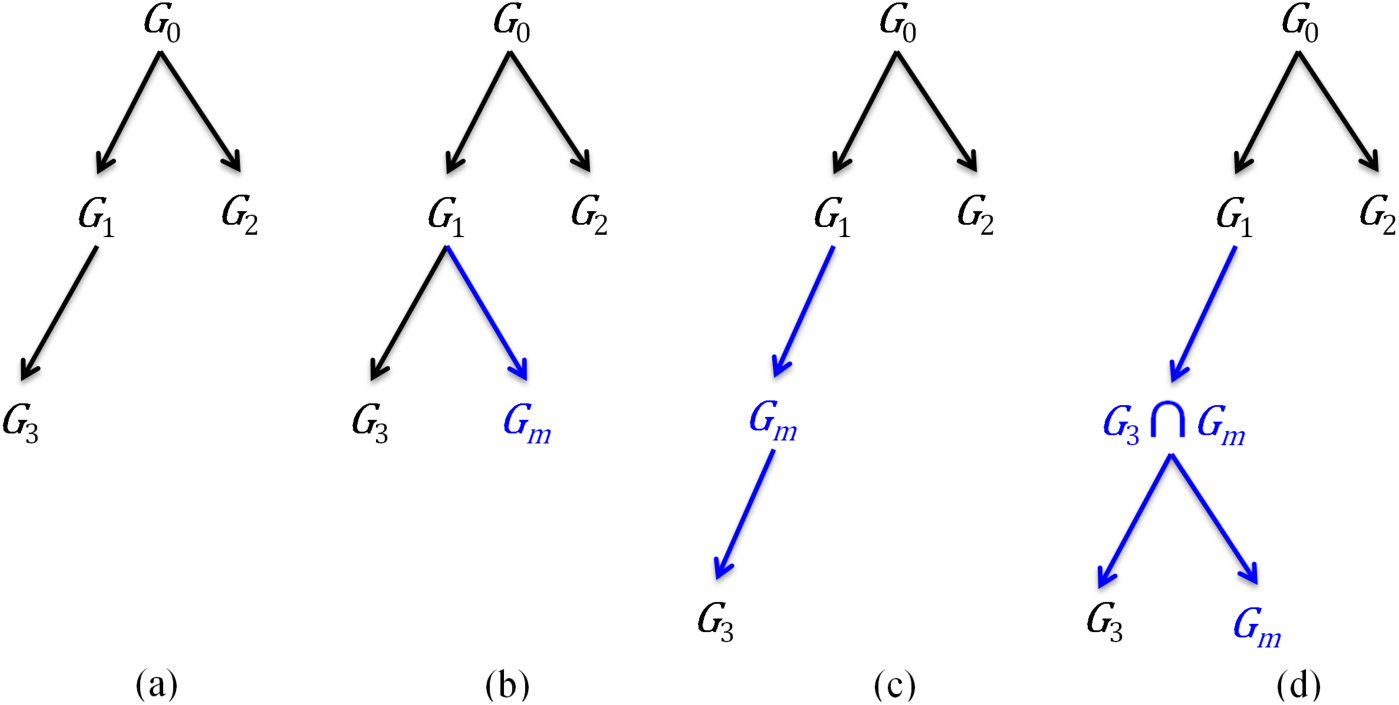
(a) An existing phylogenetic tree. (b)-(d) are three different kinds of schemes to add *G*_*m*_ to a tree. Black color indicates genomes and edges appearing in the existing tree. Blue color indicates genomes and edges that are new after adding *G*_*m*_.

After all observed tumor genomes are included in the tree, the tree grows to its full size. Because every pair of tumor genomes can be used to form an initial tree, there are 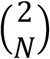 different ways of building a full-size tree, each corresponding to a unique pair of starting genomes. The algorithm constructs all 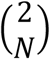 full-size trees, and selects the one with the minimum error as the final output. Algorithm 1 summarizes the entire procedure to construct a phylogenetic tree. To introduce the algorithm, we also need a measure on the similarity between two sets *X*_1_ and *X*_2_, given by

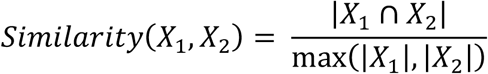

Apparently, *Similarity*(·,·) ∈ [0,1]. It is 0 when *X*_1_ and *X*_2_ do not have any overlap and is 1 when *X*_1_ is the same as *X*_2_.

#### Algorithm 1: Construction of Phylogenetic Tree

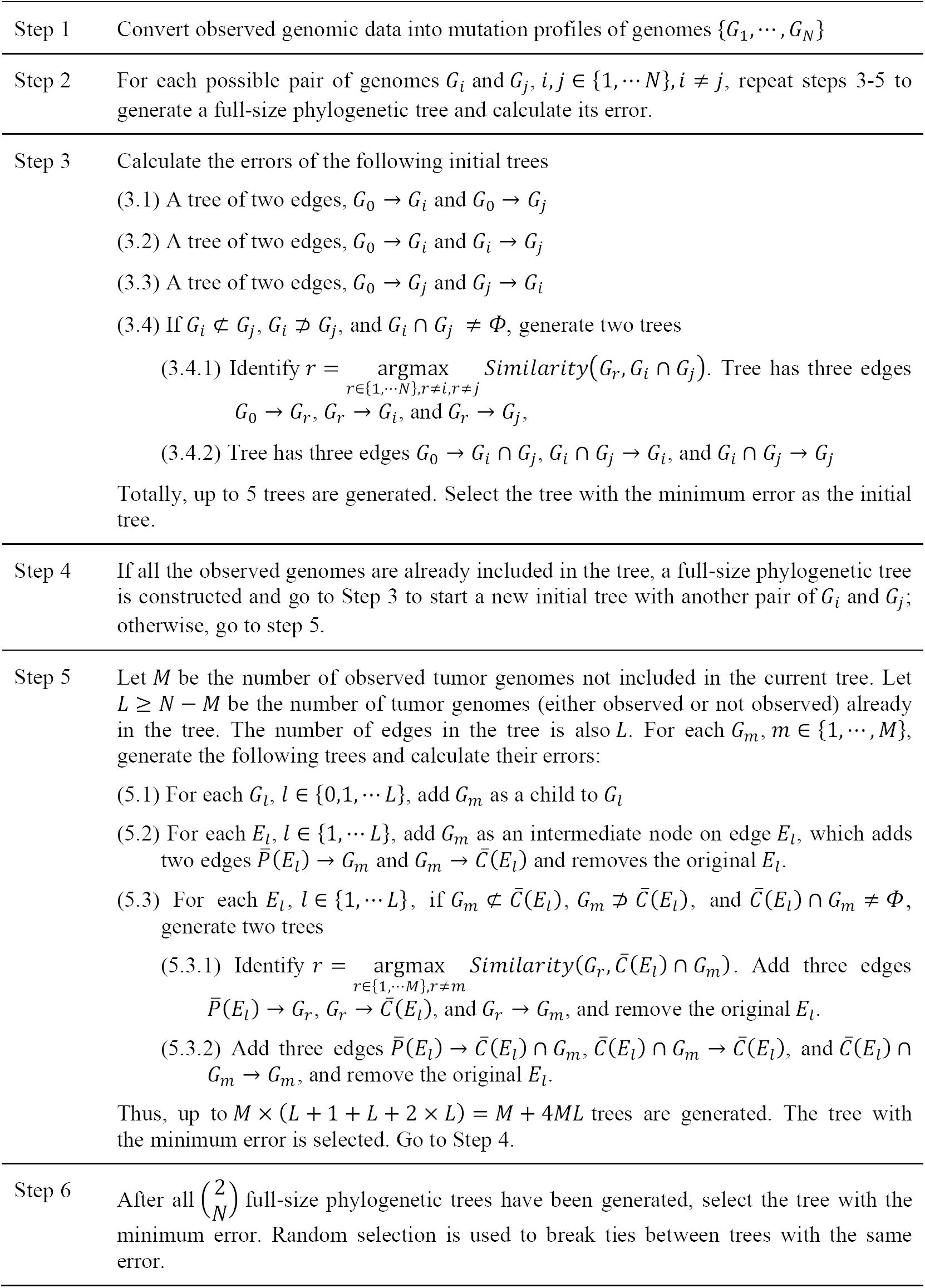

At each time when a tree grows, i.e. at Step 3 or Step 5, there may be more than one candidate tree achieving the same minimum error. Random selection can be used to break the tie. The only exception is if two candidate trees generated by Step 3.4.1 and Step 3.4.2 both give the minimum error, which is 0 in the case of the initial tree, the tree generated by Step 3.4.1 is selected, because it uses an observed genome *G*_*r*_ as the intermediate node, while Step 3.4.2 unnecessarily introduces an unobserved genome *G*_*i*_ ∩ *G*_*j*_. Similarly, if two candidate trees generated by Step 5.3.1 and Step 5.3.2 both give the minimum error in Step 5, the tree generated by Step 5.3.1 is preferred, because it does not unnecessarily introduce an unobserved genome. The purpose of including both Step 5.3.1 and Step 5.3.2 is to determine when a latent intermediate node is added, it should be an observed genome (as in Step 5.3.1) or an unobserved genome inferred by the algorithm (as in Step 5.3.2), which both belong to the scheme shown by Fig. 3d. The same consideration also supports the inclusion of both Step 3.4.1 and Step 3.4.2.

For the noise-free case where Assumptions 1 and 2 are satisfied, Theorem 2 shows that Algorithm 1 correctly identifies the phylogenetic tree.

#### Theorem 2

Assume the observed genotype data are noise-free and generated by an evolution process *T* that satisfies Assumptions 1 and 2 and thus possesses Properties 1-5. Allow that some genomes in *T* are not included in the observed genotype data. Algorithm 1 builds a phylogenetic tree 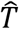 with 0 error based on the data.

The proof of Theorem 2 is given in Supplementary Information Section 4. Combining Theorem 1 and Theorem 2, on noise-free data generated from a tumor evolution process following Assumptions 1 and 2, Algorithm 1 will build a tree that has 0 error and is consistent with the underlying evolution process. Due to Step 3.4.2 and Step 5.3.2, Algorithm 1 is capable of identifying unobserved genomes that are not included in the observed data but have occurred in the evolution process. To identify an unobserved genome, two descendants from its two different child branches must be observed. These unobserved genomes may be present in the tumor but are not included in the tissue sample used for generating the data or have existed for a period of time during the evolution process and then died out.

### Edge Pruning Algorithm

In practice, all data are noisy and a tree generated by Algorithm 1 usually contains newly inferred unobserved genomes while some edges connecting with these genomes are very short, i.e. including a relatively very small number of mutation events, which may be caused by noise, because more edges with a small number of mutation events often lead to a better model fitting, i.e. a smaller error. Supplementary Information Section 5 includes an example illustrating such a situation. Motivated by this observation, we propose Algorithm 2 as a post-processing procedure to prune the phylogenetic tree generated by Algorithm 1, which overcomes potential over-fitting and prunes large trees. Algorithm 2 works with two different options. Option 1 allows the algorithm to prune the shortest edge that involves at least one unobserved genome, until the number of genomes in the tree reaches a pre-defined number. Option 2 prunes the shortest edge whose length is shorter than a pre-set threshold and that connects with at least one unobserved genome, until there is no such edge to be pruned. Notice that algorithm 2 does not remove observed genomes including the normal genome. Only unobserved genomes and their related edges may be removed. Option 1 is designed for the situation where there is a good estimation or prior knowledge about the number of genomes involved in the evolution process. In cases where such information is not available, Option 2 should be used instead.

#### Algorithm 2: Edge Pruning

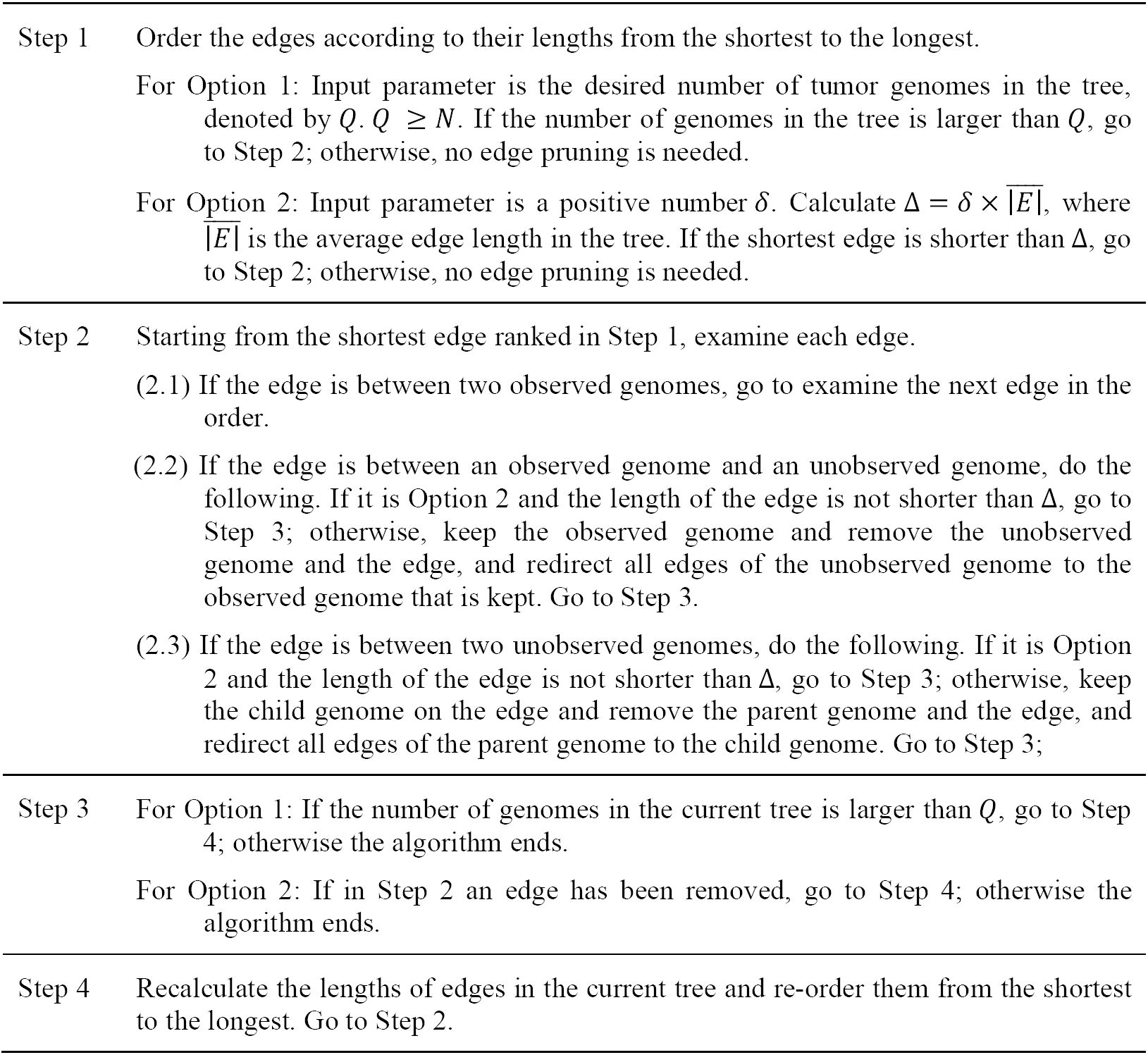

## Results

### Simulation Study

We generated simulation datasets of eight genomes according to the evolution process shown in Fig. 1. Each dataset included an SNV data matrix and a CNA data matrix (as shown in Fig. 4). Both matrices were of 200 rows and eight columns, each row representing a genomic locus and each column representing a genome. An entry at the *i*th row and *j*th column of the SNV matrix was the number of variant alleles at the *i*th locus of the *j*th genome. An entry of the CNA matrix was the copy number at a locus (indexed by its row) of a genome (indexed by its column). We divided both data matrices into five blocks. The first four blocks had the same number of rows and all rows in a block were identical with the values shown in Fig 4. These four blocks of the SNV and CNA matrices formed a portion of the genotype data that were designed to support the evolution process shown in Fig. 1. The fifth block in both the SNV and CNA matrices was composed of features with random values, which represented noise that did not support the evolutionary process and were used to test the robustness of TED. For these rows, the SNV values were uniformly sampled from {0,1,2} and the copy number values were uniformly sampled from {1,2,3}. We generated datasets with different numbers of random features (i.e. rows in the fifth block) to simulate different noise levels, ranging across 20, 40, 60, 80, 100, and 120, which constitute to 5%, 10%, 15%, 20%, 25%, and 30% of all features, respectively. At each noise level, we generated 30 simulation datasets, each with different random feature values. Therefore, we generated a total of 180 simulation datasets with noise. We generated one dataset with 0 rows in the fifth blocks, making it a noise-free dataset. Altogether, our simulation experiment included 181 datasets.

**Figure 4.**
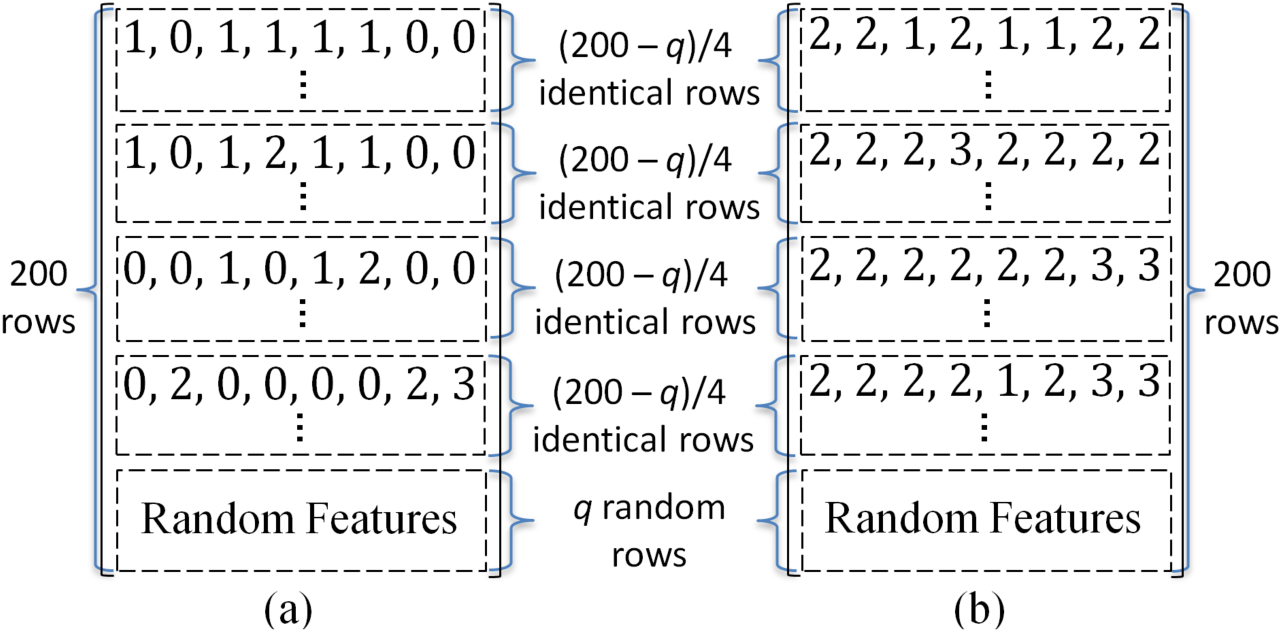
Description of the simulation data. (a) SNV data matrix. (b) CNA data matrix. Box with dashed border indicates a block in the data matrix. In both data matrices, rows in each of the first four blocks are identical. Therefore, only the first row of each block is shown.

On all the datasets including both the noise-free dataset and the noisy datasets, we ran Algorithms 1 and 2 sequentially, sending the output of Algorithm 1 as input of Algorithm 2. For Algorithm 2, both Option 1 (with *Q* = 8) and Option 2 (with *δ* = 0.5) were used for pruning edges. Option 1 with *Q* = 8 simulated the situation where there was an accurate estimation of the number of genomes involved in the evolution process. Option 2 with *δ* = 0.5 simulated the situation where there is a no such estimation or prior knowledge and a pre-set threshold on edge length (i.e. a half of the average edge length in the tree before pruning) is used instead. In cases where such information is not available, Option 2 should be used instead. Tables 3 and 4 summarize all the analysis results across the 181 datasets. At each noise level, we calculated the mean and standard deviation (s.d.) of the errors (Equation 1) associated with the estimated phylogenetic trees (see columns 2 and 3 in Tables 3 and 4). The mean and standard deviation of the consistency level (Equation 2) evaluating how accurately the estimated tree recovered the true phylogenetic tree were calculated and presented in columns 4 and 5 in the tables. For the edge pruning Option 1 (Table 3), we can see the proposed algorithms worked quite well, as the average consistency level of the estimated trees was always above 0.99 in the tested noise range. For the edge pruning Option 2, Table 4 shows that the algorithms gave a high average consistency level (≥ 0.93) when the noise level was 10% or lower. Comparing the results of the same noise level in Table 3 and Table 4, we can see for the noise level ranging from 10% to 30%, edge pruning Option 1 achieved a higher average consistency level and also a higher error than edge pruning Option 2. The reason was that edge pruning Option 2 did not prune as many edges as Option 1 did. A tree with more edges could fit the noise in data better achieving a smaller error measurement, but its topology deviated more from the truth. Also, we found that on the noise-free dataset Algorithm 1 alone (without edge pruning) constructed a phylogenetic tree identical to the ground truth shown in Fig. 1, which confirmed the validity of Theorem 2 that Algorithm 1 could construct a phylogenetic tree consistent with the underlying evolution process on noise-free data.

**Table 3.** Analysis results on simulation datasets without unobserved genome. Edges were pruned using Option 1 with *Q* = 8.

| Noise level | Error (mean) | Error (s.d.) | Consistency level (mean) | Consistency level (s.d.) |
| --- | --- | --- | --- | --- |
| 0% | 0 |  | 1 |  |
| 5% | 82.533 | 7.999 | 1 | 0 |
| 10% | 207.6 | 15.03 | 1 | 0 |
| 15% | 280.633 | 17.456 | 1 | 0 |
| 20% | 407.333 | 19.966 | 1 | 0 |
| 25% | 487.367 | 20.176 | 1 | 0 |
| 30% | 604.733 | 22.342 | 0.997 | 0.015 |

**Table 4.** Analysis results on simulation datasets without unobserved genome. Edges were pruned using Option 2 with *δ* = 0.5.

| Noise level | Error (mean) | Error (s.d.) | Consistency level (mean) | Consistency level (s.d.) |
| --- | --- | --- | --- | --- |
| 0% | 0 |  | 1 |  |
| 5% | 82.533 | 7.999 | 1 | 0 |
| 10% | 202.7 | 14.108 | 0.93 | 0.122 |
| 15% | 244.7 | 23.785 | 0.633 | 0.163 |
| 20% | 295.167 | 22.773 | 0.384 | 0.082 |
| 25% | 338.933 | 16.004 | 0.347 | 0.047 |
| 30% | 419.233 | 26.366 | 0.337 | 0.033 |

We then tested whether the proposed algorithms could identify unobserved genomes in the evolution process. For this purpose, we removed the third column (i.e. *G*_3_) of the SNV and CNA matrices in all the datasets. On each dataset, we first applied Algorithm 1 to construct a phylogenetic tree and then pruned the edges using Algorithm 2 with Option 1 (*Q* = 8) and Option 2 (*δ* = 0.5). The consistency level of the estimated tree was calculated based on the relationships between *G*_0_, *G*_1_, …, *G*_8_, including *G*_3_. Because *G*_3_ was not included in the input data, in an estimated tree we selected the unobserved genome whose mutation profile was the most similar to that of *G*_3_ to represent *G*_3_in the estimated tree, so that the consistency level could be calculated. Tables 5 and 6 summarize the results obtained using edge pruning Options 1 and 2, respectively. Algorithm 1 with the edge pruning Option 1 always achieved an average consistency level no less than 0.975 in the tested noise range. With the edge pruning Option 2, the proposed algorithms gave a high average consistency level (≥ 0.967) when the noise level was 10% or lower. Also, running Algorithm 1 alone (without edge pruning) on the noise-free dataset gave us a correct phylogenetic tree identical to Fig. 1 and also a correctly imputed *G*_3_, with all mutation events in *G*_3_ correctly inferred.

**Table 5.** Simulation results on data with *G*_3_ unobserved. Edges were pruned using Option 1 (*Q* = 8).

| Noise level | Error (mean) | Error (s.d.) | Consistency level (mean) | Consistency level (s.d.) |
| --- | --- | --- | --- | --- |
| 0% | 0 |  | 1 |  |
| 5% | 66.567 | 7.253 | 1 | 0 |
| 10% | 168.6 | 15.276 | 1 | 0 |
| 15% | 229.067 | 15.351 | 1 | 0 |
| 20% | 332.033 | 16.579 | 1 | 0 |
| 25% | 397.267 | 22.156 | 0.983 | 0.063 |
| 30% | 490.467 | 21.569 | 0.975 | 0.076 |

**Table 6.** Simulation results on data with *G*_3_ unobserved. Edges were pruned using Option 2 (δ = 0.5).

| Noise level | Error (mean) | Error (s.d.) | Consistency level (mean) | Consistency level (s.d.) |
| --- | --- | --- | --- | --- |
| 0% | 0 |  | 1 |  |
| 5% | 66.567 | 7.253 | 1 | 0 |
| 10% | 166.733 | 14.869 | 0.967 | 0.085 |
| 15% | 205.567 | 21.474 | 0.785 | 0.16 |
| 20% | 246.967 | 28.408 | 0.532 | 0.123 |
| 25% | 277.033 | 21.593 | 0.474 | 0.06 |
| 30% | 334.467 | 20.923 | 0.443 | 0.04 |

### Analysis Results of Single-Cell Sequencing Data of Essential Thrombocythemia Tumor

We analyzed a single-cell exome sequencing dataset of an essential thrombocythemia (ET) tumor without mutation in JAK2, which was published by previous literature^6,31^. JAK2 is the most commonly mutated gene among ET patients with an occurrence frequency of about 55%. The dataset included genotypes of 18 mutation sites in 18 different genes that were considered to be relevant to the disease by literatures^6,31^. The genotypes were measured for 58 single tumor cells and one normal tissue^6,31^. The normal tissue was sequenced and used as the wildtype reference, which was homozygous for the 18 mutation sites. The genotype data of these 18 mutation sites in a cell could be homozygous wildtype, heterozygous variant, and homozygous variant. Among the 18 mutation sites, 4 mutation sites in SESN2, DNAJC17, TOP1MT, and ST13 were identified with the highest likelihood of being involved with ET initiation and/or progression^6^. We performed TED analysis based on the 18 mutation sites. About 44.8% of the data entries were missing, we used K Nearest Neighbor (KNN) imputation method to generate replacements of the missing values with *K* = 5. For a missing entry in a sample, KNN imputation identified five samples closest to the given sample, whose corresponding entries were present, and then the missing entry was given the value that is most frequent among its corresponding entries in the five samples. After transferring the genotypes into mutation profiles of cells, two cells were excluded from the analysis because their mutation profiles were identical to two of other cells. Algorithm 1 was used to construct the phylogenetic tree and Algorithm 2 with Option 2 (*δ* = 0.5) was used to prune edges.

The inferred phylogenetic tree is shown in Fig. 5. It has a total error of 130 including 90 duplicated mutation events and 40 dropout mutation events. It includes 65 tumor genomes, 56 of which (i.e. c1-c56 in Fig. 5) are observed and included in the data, and the other tumor genomes (i.e. c57-c65) are inferred by the algorithm representing tumor cells involved in the evolution process but not observed in the data. The first evolution step Normal→c57 includes mutations of DNAJC17 and TOP1MT. The c61→c42 evolution step includes the mutation of SESN2. The c57→c58 evolution step includes the mutation of tumor suppressor gene ST13. Thus all these four genes most likely involved with ET initiation and/or progression^6^ mutate in the early stage of tumor evolution. Besides these four genes, an oncogene ABCB5 also mutates in the first evolution step Normal→c57 together with DNAJC17 and TOP1MT; NTRK1, a tyrosine kinase receptor that functions in a similar biological pathway as JAK2, mutates at the evolution step of c58→c59 indicating its potential role in ET progression for JAK2-negative patient.

**Figure 5.**
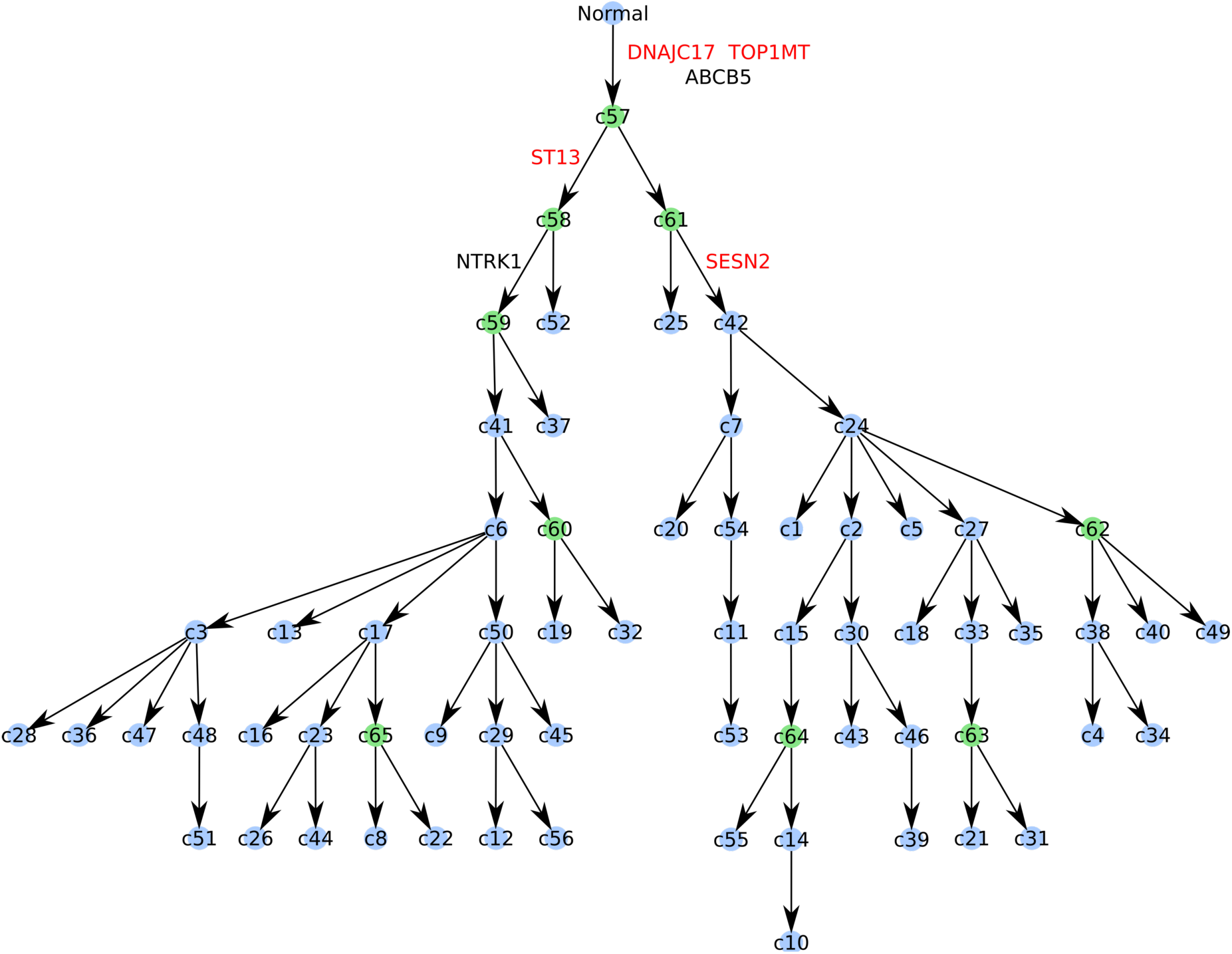
Phylogenetic tree constructed by TED on the ET tumor dataset. Cells c1-c56 (indicated by light blue) are observed genomes included in the data. Cell c57-c65 (indicated by green) are unobserved genomes inferred by the algorithm. Important genes are labeled at the evolution steps where they initially mutated. The four genes with the highest likelihood of being involved with ET initiation and/or progression6 are indicated by red.

### Analysis Results of Single-Cell Sequencing Data of Clear Cell Renal Cell Carcinoma

We applied TED on a single-cell exome sequencing dataset of a clear cell renal cell carcinoma (ccRCC), which was published by previous literature^5^. The original study sequenced 20 tumor cells and 5 normal cells, and found 3 of the tumor cells are actually normal-like. So only the 17 confirmed tumor cells were analyzed. The data included genotypes of 50 mutation sites, which could be homozygous wildtype, heterozygous variant, and homozygous variant. The normal cells were all homozygous wildtype at these mutation sites. 23.2% of the data entries were missing. We used KNN imputation with *k* = 5 to generate replacements for missing entries. Algorithm 1 was used to construct the phylogenetic tree and Algorithm 2 with Option 2 (*δ* = 0.5) was used to prune edges. Fig. 6 shows the obtained phylogenetic tree. In Fig. 6, c1-c17 are the observed genomes included in the data and c18 is an unobserved genome inferred by TED. The total error of the tree is 92 including 25 duplicated mutation events and 67 dropout mutation events. Four of the genes, i.e. SRGAP3, NIPBL, UBE4A, and SH3GL1, had been reported or predicted to contain truncating or likely functionally damaging ccRCC related mutations^5^. In the estimated phylogenetic tree, the first evolution step Normal→c10 includes the mutations of these four genes, indicating their importance of triggering tumor development. The second evolution step c10→c7 includes the mutation of RPL8, a gene whose expression level was reported to be correlated with patient response to chemotherapy^32^.

**Figure 6.**
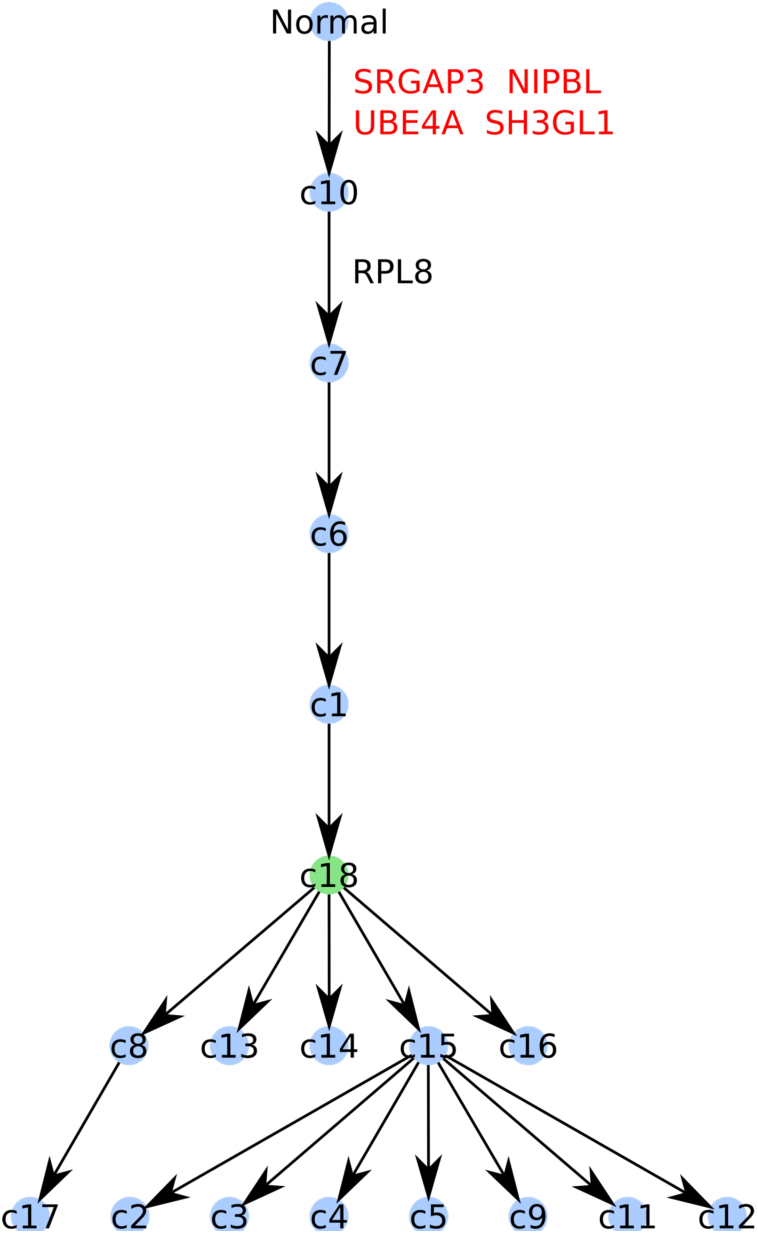
Phylogenetic tree constructed by TED on the ccRCC dataset. Cells c1-c17 (indicated by light blue) are observed genomes included in the input data. Cell c18 (indicated by green) is an unobserved genome inferred by the algorithm. Important genes are labeled at the evolution steps where they initially mutated. The four genes that have been reported or predicted to contain disease-causing ccRCC mutations are indicated by red.

### Analysis Results of Bulk Sample Subclones of Breast Invasive Carcinoma

We used TED to study a breast cancer case from The Cancer Genome Atlas (TCGA)^33^. It was a stage IIA invasive ductal carcinoma case including a primary tumor and a metastatic tumor of the same donor. We downloaded its clinical information^34^ and whole genome sequencing data ^33^ including VCF (Variant Call Format) files and BAM (Binary Alignment/Map) files. Battenberg^1^ was used to make CNA calls based on the BAM files. Both SNV data and CNA data, including 4179 SNVs and the copy numbers associated with their loci, were inputted into BayClone^16^ to infer tumor subclone genotypes. Because TCGA did not provide sequencing data of the normal tissue for this cancer case, we assumed homozygous wildtype and copy number 2 for all loci as the normal genotype. BayClone first performed tumor purity analysis and then within the tumor portion it identified three tumor subclones, indicated by subclones 1, 2, and 3 in Fig. 7. BayClone inferred the SNVs and CNAs of the three subclones, and their cellular proportions in the primary tumor and metastatic tumor (see Fig. 7). We then used TED to study the evolution process of the subclones based on their SNVs and CNAs. Algorithm 1 constructed the phylogenetic tree and Algorithm 2 with Option 2 (*δ* = 0.5) was used to prune edges. TED analysis gave a phylogenetic tree consisting of four tumor subclones (see Fig. 7), in which subclone 4 is an unobserved genome inferred by TED. For SNVs and CNAs on each evolution step, we selected the mutations in the Coding DNA Sequence (CDS), which were labeled as CDS-SNVs and CDS-CNAs, respectively, and then used the Bioconductor package EGSEA^35^ to carry out Gene Set Enrichment Analyses (GSEA) for the genes that harbored these mutations. GSEA was done separately for the genes with CDS-SNVs and for the genes with CDS-CNAs. The gene sets used in the analyses included the Molecular Signatures Database (MSigDB) v5.0^36^, KEGG pathways^37^, and GeneSetDB^38^. A gene set was deemed as statistically significant, if its adjusted p-value is no larger than 0.05.

**Figure 7.**
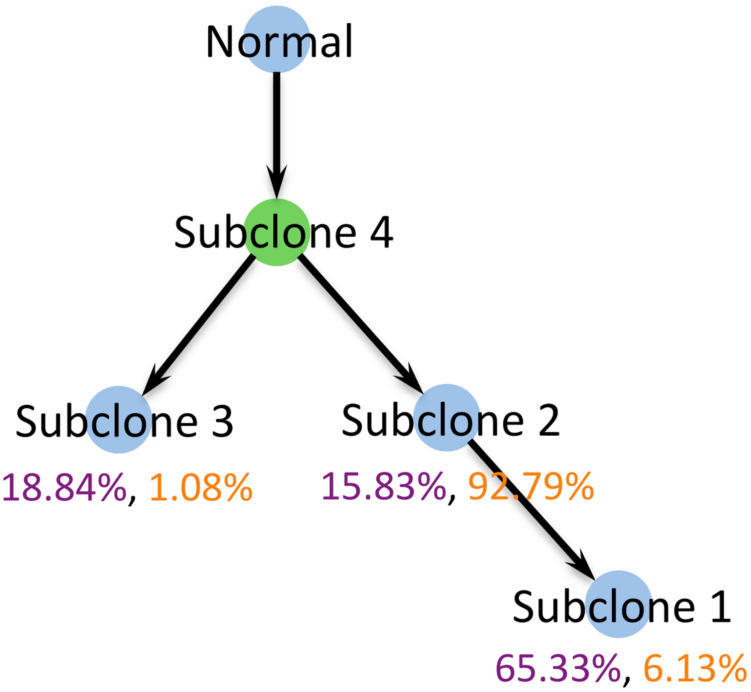
Phylogenetic tree constructed by TED for a TCGA breast invasive carcinoma case based on statistically inferred subclones (indicated by light blue). TED infers an unobserved genome that is subclone 4 indicated by green. The numbers under subclones 1, 2, and 3 are their statistically inferred cellular proportions in the tumors. Purple indicates the cellular proportion in the primary tumor and orange indicates the cellular proportion in the metastatic tumor.

GSEA shows that RhoA regulation related pathways that regulate cell shape, attachment, and motility are statistically significantly enriched in the CDS-SNV genes on the normal → subclone 4 edge, including DLC1, an important gene in the RhoA regulation related pathways that has been reported with the potential of suppressing breast cancer bone metastasis and whose mutation may lead to metastasis^39^. The RhoA regulation related pathways are also significantly enriched in the CDS-CNA genes on the edge of subclone 4 → subclone 2, including ARHGAP45 (Rho GTPase activating protein 45) that has been reported to act as a RhoGAP to regulate GTPase activity, cytoskeletal remodeling and cell spreading^40^. The copy number of ARHGAP45 is amplified in this evolution step, leading to more metastasis potential of subclone 2. Also, in this evolution step, SNV mutations occur to CBFA2T3, a tumor suppressor gene of breast cancer^41^. On the edge of subclone 4 → subclone 3, another RhoA regulation related gene, ARHGEF7 (Rho guanine nucleotide exchange factor 7), has SNV mutations that may potentially weak its inhibition of cell migration and proliferation in breast cancer^42^. Cytoskeletal and extracellular-matrix related genes are significantly enriched in the CDS-SNV genes and CDS-CNA genes on the edge of subclone 2 → subclone 1, which may change the cell motility and microenvironment adaptability of tumor cells, respectively, and thus affect the growth of subclones. Subclone 1 has the largest cell population (65.33%) in the primary tumor, but its population is very small (6.13%) in the metastasis tumor. Subclone 2 constitutes to a minor component (15.83%) of the primary tumor but it is dominant (92.79%) in the metastasis tumor. Subclone 1 and subclone 2 may migrate to the metastatic site together through polyclonal seeding^43^, but subclone 2 is much more adaptive to the metastatic site than subclone 1 that prospers at the primary site. Some of the mutated genes on the subclone 2 → subclone 1 edge have been reported important in cancer, such as IGFALS (insulin like growth factor binding protein acid labile subunit) that has CDS-SNV mutations and CLIP1 (CAP-Gly domain containing linker protein 1) that has copy number amplifications in its CDS. IGFALS is a serum protein that binds insulin-like growth factors, increasing their half-life and their vascular localization. It has been reported to be a tumor suppressor in hepatocellular carcinomas^44^. CLIP1 links endocytic vesicles to microtubules, and was found to be related to invasive breast cancer^45^.

## Discussion

We have developed TED, a novel algorithm for constructing phylogenetic tree from a new perspective that has not been explored before by other phylogenetic tree methods. TED constructs a phylogenetic tree by reducing the total number of duplicated mutations and dropout mutations in the evolution process and can be applied to genotype data of either single tumor cells or tumor subclones. The validity of TED was shown through newly proved theorems. We tested TED on simulation datasets. If the true number of distinct tumor genomes involved in the evolution process was known a priori, TED could achieve a very high performance (average consistency level ≥ 0.997), when up to 30% of the features were purely noise. If the number of tumor genomes was not known, TED could achieve an average consistency level ≥ 0.93, when 10% or less of the features were noise. The simulation study also showed if a tumor genome was not observed and not included in the input data, TED could still achieve a high performance for recovering the entire evolution process including the unobserved tumor genome. We then applied TED on two single-cell exome sequencing datasets, an ET tumor case and a ccRCC case. In both cases, we found genes that were reported (by previous works) important for cancer initialization and development mutated in the early steps of the recovered phylogenetic tree. We also applied TED on the mutation profiles of statistically inferred subclones for a TCGA breast cancer case. The constructed phylogenetic tree showed meaningful insights on the evolution and prevalence of tumor subclones.

TED is different from existing phylogenetic tree construction methods. It is specifically designed for tumor evolution study and emphasizes on correctly recovering the ancestor-descendant relationships between tumor genomes, i.e. the topology of evolution process. TED includes not only a phylogenetic tree construction algorithm but also an edge pruning algorithm, which allows the evaluation of TED performance by comparing the edge-pruned tree directly with the ground truth evolution tree in terms of the tree topology. Most other phylogenetic tree algorithms do not have a such edge pruning function and usually include in their outputs a significant number of intermediate nodes that do not correspond to any observed tumor genome in the input data. They usually take all the input genomes as the leaf nodes in the tree plot, which destroys all the ancestor-descendant relationships between genomes making it infeasible to compare their outputs directly with the ground truth evolution tree for topology evaluation. Thus, we have not included other methods in our simulation study for performance comparison with TED.

Some existing methods for tumor heterogeneity study integrate subclone inference with evolution tree construction for analyzing DNA sequencing data of bulk sample^30,46^. TED has been developed for the sole purpose of revealing the topology of a tumor evolution process. It constructs a phylogenetic tree based on known normal and tumor genomes and does not infer tumor subclone genotypes based on DNA sequencing data. Thus, its applications can be different from the tumor heterogeneity methods. For example, we have applied TED for single-cell sequencing data analysis.

The proof of Theorem 2 does not utilize any information about which two genomes are selected to generate the initial tree and in what order the other genomes are added to the tree. This indicates, for noise-free data following Assumptions 1 and 2, Algorithm 1 can be simplified by randomly selecting two genomes to construct an initial tree and randomly selecting a candidate genome to add to the tree in the later steps. There is no need to exhaustively search among all initial genome pairs and among all candidate genomes to be added in the later steps. The full-size tree generated will be error-free and consistent with the underlying evolution process.

The TED algorithm minimizes the error of phylogenetic tree in a stepwise manner. When generating an initial tree or adding a genome to an existing tree, all possible candidate genomes and all possible schemes to generate the tree are considered to identify the tree with the minimum error at each time. Although this stepwise minimization approach does not guarantee a global optimization, our simulation study shows that TED achieved a good estimation accuracy of the evolution process even when the input data contain significant noise and our applications on real data have generated biologically plausible and interesting results.

The error function of TED includes two types of errors, i.e. the number of duplicated mutations (Type I error) and the number of dropout mutations (Type II error). Currently, they are equally weighted in the error function. A variant of the TED algorithm can be made by adding a multiplication factor to either Type I or Type II error, which gives different weights to different types of errors. Furthermore, different kinds of mutations can have different weights too. The modified error function can penalize the error types or mutation types with heavy weights, thus reduce their numbers more severely. Such an algorithm may have different performance compared to the current TED algorithm and may be applicable in different data analysis situations, which can be investigated in future research. Notice that because all the weights are positive, Theorems 1 and 2 will still hold for these variant algorithms, which means Algorithm 1 with a variant error function can still construct a phylogenetic tree consistent with the underlying evolution process on ideal, noise-free data.

We implemented TED using the R programming language and provide it as open-source, free-available software at http://compgenome.org/ted/

## Author Contributions

Y.Z. and Y.J. conceived the main idea and participated in all aspects of the project. Y.Z. developed the TED algorithm, derived the supporting theory, and performed the analyses. S.S., L.W., and S.Y. participated in TCGA data analysis and annotated the analysis results. All authors edited the manuscript.

