## Supplementary Information for "Tumor Evolution Decoder (TED): Unveiling Tumor Evolution Based on Mutation Profiles of Subclones or Single Cells"

Yitan Zhu[*1], Subhajit Sengupta[1], Lin Wei[1], Shengjie Yang[1], Yuan Ji[*1,2]

1. Program of Computational Genomics & Medicine, NorthShore University HealthSystem, Evanston, Illinois, USA
2. Department of Public Health Sciences, The University of Chicago, Chicago, Illinois, USA

* Corresponding author

***Corresponding Authors***

Yitan Zhu
1001 University Pl, Evanston, IL 60201


Yuan Ji
1001 University Pl, Evanston, IL 60201


#### Section 1 Proof of Property 4

Apparently, $G_i$ and $G_j$ must have one or more common ancestors in $T$, because at least $G_0$ is a common ancestor. Their common ancestors must be a group of nested sets, so there is one common ancestor that includes all other common ancestors as subsets. Let $G_c$ be the largest common ancestor. $G_c \in \{G_0, G_1, G_2, \cdots G_N\}$. $G_c$ is also the closest to $G_i$ and $G_j$ among all common ancestors, because all other common ancestors are its subsets. Because $G_c \subset G_i$ and $G_c \subset G_j$, $G_c \subseteq G_i \cap G_j$. Suppose that there is a mutation event $u \in G_c$ and $u \notin G_i \cap G_j$. $u$ must occur on both the path from $G_c$ to $G_i$ and the path from $G_c$ to $G_j$, which conflicts with Assumption 1. So such $u$ can not exist, and $G_c = G_i \cap G_j$. Thus, $G_i \cap G_j$ is the closest to $G_i$ and $G_j$ and also the largest among all their common ancestors.

#### Section 2 Proof of Property 5

If $G_i \subset G_j$, then $G_i \cap G_j = G_i \in \{G_0, G_1, G_2, \cdots G_N\}$. If $G_j \subset G_i$, then $G_i \cap G_j = G_j \in \{G_0, G_1, G_2, \cdots G_N\}$. If $G_i \not\subset G_j$ and $G_i \not\supset G_j$, because $G_i \neq G_j$, then $G_i \nsubseteq G_j$ and $G_i \nsupseteq G_j$. According to property 4, $G_i \cap G_j$ is a common ancestor of $G_i$ and $G_j$ in $T$. So in all of the above three possible cases, $G_i \cap G_j \in \{G_0, G_1, G_2, \cdots G_N\}$.

#### Section 3 Proof of Theorem 1

If $\hat{T}$ is not consistent with $T$, there must be at least one pair-wise relationship being changed. Let them be $G_i$ and $G_j$ in $T$. The change can be one of the three following cases.

(1) $G_i$ and $G_j$ has an ancestor-descendant relationship in $T$. Without loss of generality, let $G_i$ be an ancestor of $G_j$. In $\hat{T}$, $G_j$ becomes an ancestor of $G_i$. Then, there are back mutations that are Type II errors.

(2) $G_i$ and $G_j$ has an ancestor-descendant relationship in $T$. Let $G_i$ be an ancestor of $G_j$. In $\hat{T}$, $G_i$ and $G_j$ do not have an ancestor-descendant relationship, which means they are no longer both in a path from $G_0$ to a leaf node. In such a case, consider two paths in $\hat{T}$, which are from the closest common ancestor of $G_i$ and $G_j$ to $G_i$ and $G_j$. There must be mutations happening on both paths that cause Type I errors.

(3) In $T$, $G_i$ and $G_j$ do not have an ancestor-descendant relationship, which indicates $G_i - G_j \neq \Phi$ and $G_j - G_i \neq \Phi$. But in $\hat{T}$, they have an ancestor-descendant relationship. Let $G_i$ be an ancestor of $G_j$. Then, there are dropout mutations that are included in $G_i$ but not in $G_j$ and cause Type II errors.

So if $error(\hat{T}) = 0$, $\hat{T}$ is consistent with $T$. Without loss of generality, let the genomes in $\hat{T}$ be $G_0, G_1, \cdots, G_n, n \leq N$. Suppose that $\hat{T}$ is not closed under intersection. Then there exist two genomes $G_i$ and $G_j$, $i, j \in \{1, \cdots, n\}, G_i \cap G_j \notin \{G_0, G_1, \cdots, G_n\}$. $G_i$ and $G_j$ must have at least one common ancestor in $\hat{T}$, because at least $G_0$ is their common ancestors. The common ancestors of $G_i$ and $G_j$ must be a group of nested sets. Let $G_k$ be the largest of them, where $k \in \{0,1,\cdots,n\}, k \neq i, k \neq j$. $G_i \cap G_j$ must be the same as $G_k$, because otherwise the path from $G_k$ to $G_i$ and the path from $G_k$ to $G_j$ will share at least one mutations that cause Type I error. So $\hat{T}$ must be closed under intersection.

### Section 4 Proof of Theorem 2

Actually, we can prove starting from an initial tree constructed using any pair of genomes, Algorithm 1 Steps 4 and 5 will build a full-size phylogenetic tree with 0 error.

Algorithm 1 Step 3 constructs an initial phylogenetic tree with the normal genome $G_0$ and two tumor genomes (let them be $G_1$ and $G_2$). We denote the initial tree by $\hat{T}_{ini}$. There are three possible relationships between $G_1$ and $G_2$.

(1) $G_1$ and $G_2$ are from two different lineages in $T$, so $G_1 \cap G_2 = G_0$. The tree generated by Step 3.1 will be selected as $\hat{T}_{ini}$, which has 0 error.

(2) $G_1$ and $G_2$ are in the same lineage and have an ancestor-descendant relationship, i.e. either $G_1 \subset G_2$ or $G_2 \subset G_1$. A tree generated in either Step 3.2 or Step 3.3 will be selected as $\hat{T}_{ini}$, which has 0 error.

(3) $G_1$ and $G_2$ are in the same lineage, but do not have an ancestor-descendant relationship. In this case, $G_1 \cap G_2 \neq \Phi, G_1 \not\subseteq G_2$, and $G_1 \not\supseteq G_2$. A tree generated in Step 3.4 will be selected as $\hat{T}_{ini}$ and it has 0 error.

So in all three cases, $\hat{T}_{ini}$ has 0 error.

Then, suppose we have constructed a phylogenetic tree $\hat{T}$ that includes $G_0, G_1, \cdots, G_n$ and that $\hat{T}$ has 0 error, which indicates $\hat{T}$ is closed under intersection and consistent with $T$. Consider adding $G_m$ to $\hat{T}$ to generate $\hat{T}_{next}$. There are three possible relationships between $G_m$ and $G_0, G_1, \cdots, G_n$.

(1) $G_m$ is not an ancestor of any of $G_1, \cdots, G_n$ in $T$ and $G_0, G_1, \cdots, G_n, G_m$ are closed under intersection. This means $\forall i \in \{1, \cdots, n\}, G_i \not\supset G_m$ and $\forall i \in \{0,1,\cdots,n\}, G_i \cap G_m \in \{G_0, G_1, \cdots, G_n\}$. $A(G_m)$, the set of all ancestors of $G_m$ in $T$, must not be empty, because it contains at least $G_0$ that appears in both $T$ and $\hat{T}$. $A(G_m)$ must be a group of nested sets. Let $G_{i^*}$ be the largest set in $A(G_m)$ that is already included in $\hat{T}$. Algorithm 1 Step 5.1 can add $G_m$ as a child node of $G_{i^*}$ to generate $\hat{T}_{next}$. Apparently, $\hat{T}_{next}$ does not have any Type II error, because there is no back/dropout mutation on the newly added edge $G_{i^*} \rightarrow G_m$. Suppose $\hat{T}_{next}$ has Type I error, which must be caused by some shared mutation event between $G_{i^*} \rightarrow G_m$ and some edge already included in $\hat{T}$. Let $u$ be such a mutation event. Then, the following two conditions must hold.

(1.a) $\forall j \in \{1, \cdots, n\}$ and $G_j \in A(G_m), u \notin G_j$, because $u \notin G_{i^*}$, which is the largest set in $A(G_m)$.

(1.b) Therefore, $\exists k \in \{0,1,\cdots,n\}, G_k \notin A(G_m)$ and $u \in G_k$.

So $u \in G_k \cap G_m \in \{G_0, G_1, \cdots, G_n\}$, because $G_0, G_1, \cdots, G_n, G_m$ are closed under intersection. Apparently, $G_k \cap G_m \in A(G_m)$ and $u \in G_k \cap G_m$, which conflicts with (1.a). So $\hat{T}_{next}$ does not have any Type I error.

(2) $G_m$ is an ancestor of some genome among $G_1, \cdots G_n$ in $T$, which indicates $\exists i, j \in \{0,1, \cdots, n\}, G_i \subset G_m \subset G_j$. All genomes on the path from $G_i$ to $G_j$ in $\hat{T}$ must also be either an ancestor or a descendent of $G_m$ in $T$, because $\hat{T}$ is consistent with $T$. Among them, let $G_{j^*}$, $j^* \in \{0,1, \cdots, n\}$, be the smallest descendent of $G_m$ that is already in $\hat{T}$ and $G_{i^*}$, $i^* \in \{0,1, \cdots, n\}$, be the largest ancestor of $G_m$ that is already in $\hat{T}$. The edge $G_{i^*} \rightarrow G_{j^*}$ must exist in $\hat{T}$. Step 5.2 can add $G_m$ as an intermediate node on this edge and generate a 0-error $\hat{T}_{next}$.

(3) $G_m$ is not an ancestor of any of $G_1, \cdots, G_n$ in $T$, and $G_0, G_1, \cdots, G_n, G_m$ are not closed under intersection. This means $\forall i \in \{1, \cdots, n\}, G_i \not\supset G_m$, and $\exists i \in \{0,1, \cdots, n\}$, $G_i \cap G_m \notin \{G_0, G_1, \cdots, G_n, G_m\}$. Let $G_{i^*}$, $i^* \in \{0,1, \cdots, n\}$ be the largest ancestor of $G_m$ that is included in $\hat{T}$. Apparently, in $\hat{T}$ only the descendants of $G_{i^*}$ can have an intersection with $G_m$ that falls out of $\{G_0, G_1, \cdots, G_n, G_m\}$, because $\forall k \in \{1, \cdots, n\}$ and $G_k \notin D(G_{i^*})$, $G_k \cap G_m = G_k \cap G_{i^*} \in \{G_0, G_1, \cdots, G_n, G_m\}$. Consider a child of $G_{i^*}$ in $\hat{T}$ denoted by $G_j$, $G_m \cap G_j$ is either $G_{i^*}$ or not included in $\{G_0, G_1, \cdots, G_n, G_m\}$. If $G_m \cap G_j = G_{i^*}$, which means $G_{i^*}$ is the closest common ancestor to $G_m$ and $G_j$ in $T$, then $\forall s \in \{1, \cdots, n\}$ and $G_s \in D(G_j)$, $G_m \cap G_s = G_{i^*} \in \{G_0, G_1, \cdots, G_n, G_m\}$. So there must be at least one child of $G_{i^*}$ whose intersection with $G_m$ is not included in $\{G_0, G_1, \cdots, G_n, G_m\}$.

Suppose that $\exists j_1, j_2 \in \{1, \cdots, n\}$, both $G_{j_1}$ and $G_{j_2}$ are children of $G_{i^*}$ in $\hat{T}$ and that both $G_m \cap G_{j_1}$ and $G_m \cap G_{j_2}$ are not in $\{G_0, G_1, \cdots, G_n, G_m\}$. $G_m \cap G_{j_1}$ and $G_m \cap G_{j_2}$ can not be the same; otherwise $G_m \cap G_{j_1} = G_m \cap G_{j_2} \supset G_{i^*} \Longrightarrow G_{j_1} \cap G_{j_2} \supset G_{i^*}$, giving the error of duplicated mutations on the edge $G_{i^*} \rightarrow G_{j_1}$ and the edge $G_{i^*} \rightarrow G_{j_2}$ in $\hat{T}$. So $G_m \cap G_{j_1}$ and $G_m \cap G_{j_2}$ are different. Then, in $T$ there are two paths from $G_{i^*}$ to $G_m$, one through $G_m \cap G_{j_1}$ and the other through $G_m \cap G_{j_2}$, which also cause duplicated mutations in $T$. Thus, there must be one and only one child of $G_{i^*}$ in $\hat{T}$ (denoted by $G_{j^*}$) that gives $G_m \cap G_{j^*} \notin \{G_0, G_1, \cdots, G_n, G_m\}$.

Algorithm 1 Step 5.3 will add two genomes, i.e. $G_m \cap G_{j^*}$ and $G_m$, to $\hat{T}$ as illustrated by Fig. 3d. Apparently, the resulted $\hat{T}_{next}$ does not have any Type II error. Because $G_m \cap G_{j^*}$ is the closest common ancestor to $G_m$ and $G_{j^*}$ in $T$, the edge $G_m \cap G_{j^*} \rightarrow G_m$ will not share any mutation with other edges in $\hat{T}_{next}$. So $\hat{T}_{next}$ does not have any Type I error neither.

In all of the three possible cases, Algorithm 1 Step 5 will always generate an error-free $\hat{T}_{next}$. Thus, when Algorithm 1 ends, the full-size phylogenetic tree must have 0 error, and thus is consistent with $T$.

#### Section 5 An Example Of Edge Pruning

We pick one of the simulation datasets used for performance evaluation to illustrate the edge pruning effect. It is a dataset of 5% noise level, i.e. 20 out of the 400 mutation features are random noise. Fig. S1b and Fig. S1c show the phylogenetic trees before and after edge pruning, respectively, where the true tree is given in Fig. S1a. In this case, both Options of Algorithm 2 give the same pruned tree that is identical to the true tree. Option 1 is set to keep 8 tumor genomes and Option 2 is set to remove edges whose lengths are shorter than 50% of the average edge length in the tree before edge pruning starts.

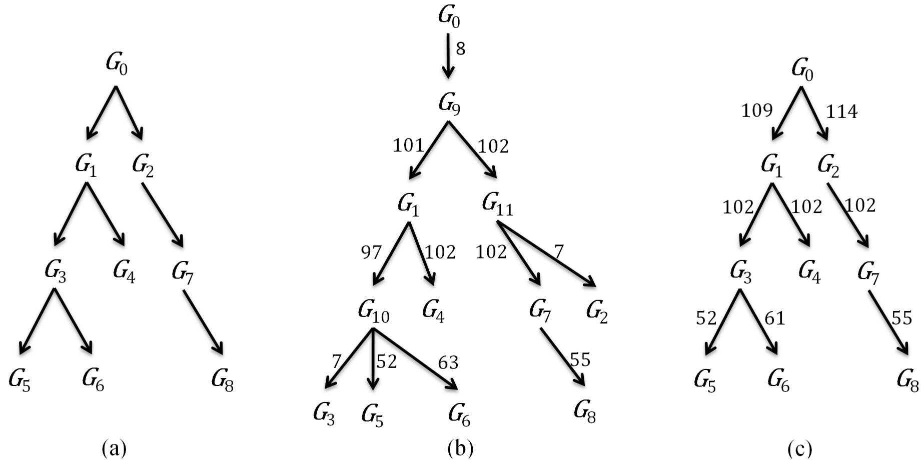


**Figure S1** An illustration of using Algorithm 2 to prune noisy edges in a tree constructed by Algorithm 1 (a) The evolution process used for generating the simulation data. (b) The estimated phylogenetic tree constructed by Algorithm 1 without pruning. The numeric values on the edges are the edge lengths. (c) The estimated phylogenetic tree obtained after pruning edges using Algorithm 2. Both Option 1 and Option 2 give the same pruned tree, which is consistent with the ground truth (a).
